# A hybrid bioprinting-electrospinning platform integrating nanofibers and mesenchymal cell spheroids for customizable wound healing dressings

**DOI:** 10.1101/2024.09.14.613065

**Authors:** Seyede Atefe Hosseini, Viktoria Planz, Ernst H. K. Stelzer, Maike Windbergs, Francesco Pampaloni

## Abstract

We introduce a platform for the fabrication of customizable wound healing dressing. The platform integrates electrospun nanofibers, bioprinted hydrogels, and cellular spheroids into hierarchical, fiber-reinforced hybrid constructs. The construct leverages the mechanical strength of polycaprolactone (PCL) nanofibers and the ECM-like properties of GelMA/PEGDA hydrogel. These materials support the incorporation of bone marrow-derived mesenchymal stem cell (BM-hMSC) spheroids, which act as a supportive “cell niche,” enhancing the viability of the hMSC during and after bioprinting, and facilitating their spreading across the construct during the maturation phase. The characterization of the hybrid constructs demonstrated strong structural integrity and enhanced mechanical properties, making them well-suited for clinical wound dressing applications. *In vitro* assays, including live/dead staining, MTT assays, and scratch assays, revealed increased cell attachment, proliferation, and migration. The spheroids maintained their viability over extended periods, significantly contributing to wound closure in the scratch assay. This innovative approach, which combines electrospinning and light-based bioprinting, offers a promising strategy for the development of customizable wound dressings that closely adapt to the complex architecture of human skin. The bioprinting approach allows for the creation of tailored geometries for specific clinical requirements. Future research will focus on optimizing scaffold design and conducting long-term *in vivo* studies to validate the platform’s clinical potential.

## Introduction

In recent years, in vitro skin scaffolds and models have been developed and utilized for various clinical and research purposes, including skin grafting for wound or burn healing, wound dressing, and studies on skin physiology and pathophysiology (1, 2). Among skin damages, diabetic wounds are particularly challenging, often resisting healing due to a complex interplay of impaired blood flow, neuropathy, and chronic inflammation (3). These wounds are a major complication of diabetes mellitus, increasing the risk of clinical infection, disability, and even death. In 2019, approximately 463 million adults globally had diabetes, with this number projected to rise to 578 million by 2030 and 700 million by 2045 (4). Hence, it is crucial to develop wound dressings that not only act as physical barriers to prevent further damage and infection but also promote wound closure, aiding in faster healing and reduced scar formation. Among various dressing materials, hydrogel models, particularly lattice models, are the most widely used in vitro skin models in research. These models typically employ cells, biomaterials, and proteins (5).

Despite the biocompatibility of hydrogel models and their ability to facilitate cell and drug loading, they suffer from low extracellular matrix (ECM) neosynthesis and poor mechanical strength and stability (6–8). Therefore, there is a pressing need to develop more functional scaffolds that can effectively repair complex wounds. This task is challenging due to the intricate and heterogeneous nature of human skin (9, 10).

One alternative is to use porous materials or scaffolds, which allow for cell migration, attachment, and ECM neosynthesis, thereby creating a dynamic, complex, and organized dermal compartment and a microenvironment that closely mimics in vivo skin (11, 12). Porous scaffolds can be made from biological materials such as collagen, elastin, or hyaluronic acid, either individually or in combination, or from synthetic materials such as polystyrene or poly(ε-caprolactone) (PCL) (13). Techniques for creating these scaffolds include electrospinning, 3D bioprinting, self-assembly, phase separation, particulate leaching, melt extrusion, and gas foaming. Scaffold architecture aims to closely replicate dermal organization, though many techniques fall short of this goal (5).

Current porous scaffold techniques have a limited ability to faithfully replicate the ultrastructural architecture of the skin, likely due to manufacturing constraints (14). To address the need for a functional skin construct and the limitations of existing porous systems, we combined electrospinning and bioprinting to create a hybrid gelatin methacrylate (GelMA) and Polyethylene Glycol Diacrylate (PEGDA) scaffold/PCL mesh construct. In this construct, the PCL membrane serves as the outer layer exposed to the external environment (like the epidermis), while the printed scaffold provides an optimal open-pore structure for the dermis. The PCL membrane’s porosity allows for gas exchange while limiting cell infiltration, with high porosity and small pore size, yielding a large surface area and a structure capable of mimicking epidermal features. These scaffolds have good biocompatibility and long-term biodegradability, with excellent, tailorable mechanical properties (13). Research has shown that bioprinted constructs enhance cell attachment, growth, and differentiation (15), making them a promising component in our innovative scaffold/membrane construct.

In this paper, we present evidence that combining electrospinning and bioprinting allows for the rapid development of a skin scaffold that supports superior attachment and proliferation of human bone marrow mesenchymal stem cell (BM-hMSC) spheroids. The embedded BM-hMSC spheroids work as “cell reservoir” from which the BM-hMSC migrate, proliferate, and progressively populate the construct. Using BM-hMSC spheroids enhances cell viability during seeding and bioprinting as well as during the time required for the construct’s maturation. Thus, this research showcases the creation of skin constructs with minimal initial ECM content and BM-hMSC spheroids instead of single BM-hMSCs.

Our study demonstrates how the chosen scaffold designs influence ECM organization, diabetic wound healing, and mechanical properties.

These findings position bicomponent scaffolds as a promising advancement for rapidly creating diverse skin scaffolds and models with tunable geometry and mechanical properties. Such models have the potential to significantly advance skin-related knowledge, facilitate evaluation studies, and possibly serve as skin grafts.

## Results

### Characterization of electrospun PCL nanofiber scaffolds

Single-component non-woven electrospun nanofiber scaffolds were fabricated by using PCL, a mechanically stable, synthetic polymer exhibiting slow-degrading properties in an aqueous physiological environment. The scaffolds revealed a beadless fiber network with a smooth surface as visualized by SEM micrographs (**Figure 1a**). However, a few considerably smaller fibers (< 1.0 µm) were apparent. In total, PCL scaffolds exhibited fiber diameters in the low micrometer range from 0.8 to 5.0 µm with an average of 2.89 µm ± 0.98 µm (**Figure 1b**). The percentage fiber diameter distribution with Gaussian fitting is demonstrated in **Figure 1c**. The average thickness of the fabricated scaffolds was characterized as 223.16 µm ± 18.85 µm (**Figure 1d**).

**Figure 1.**
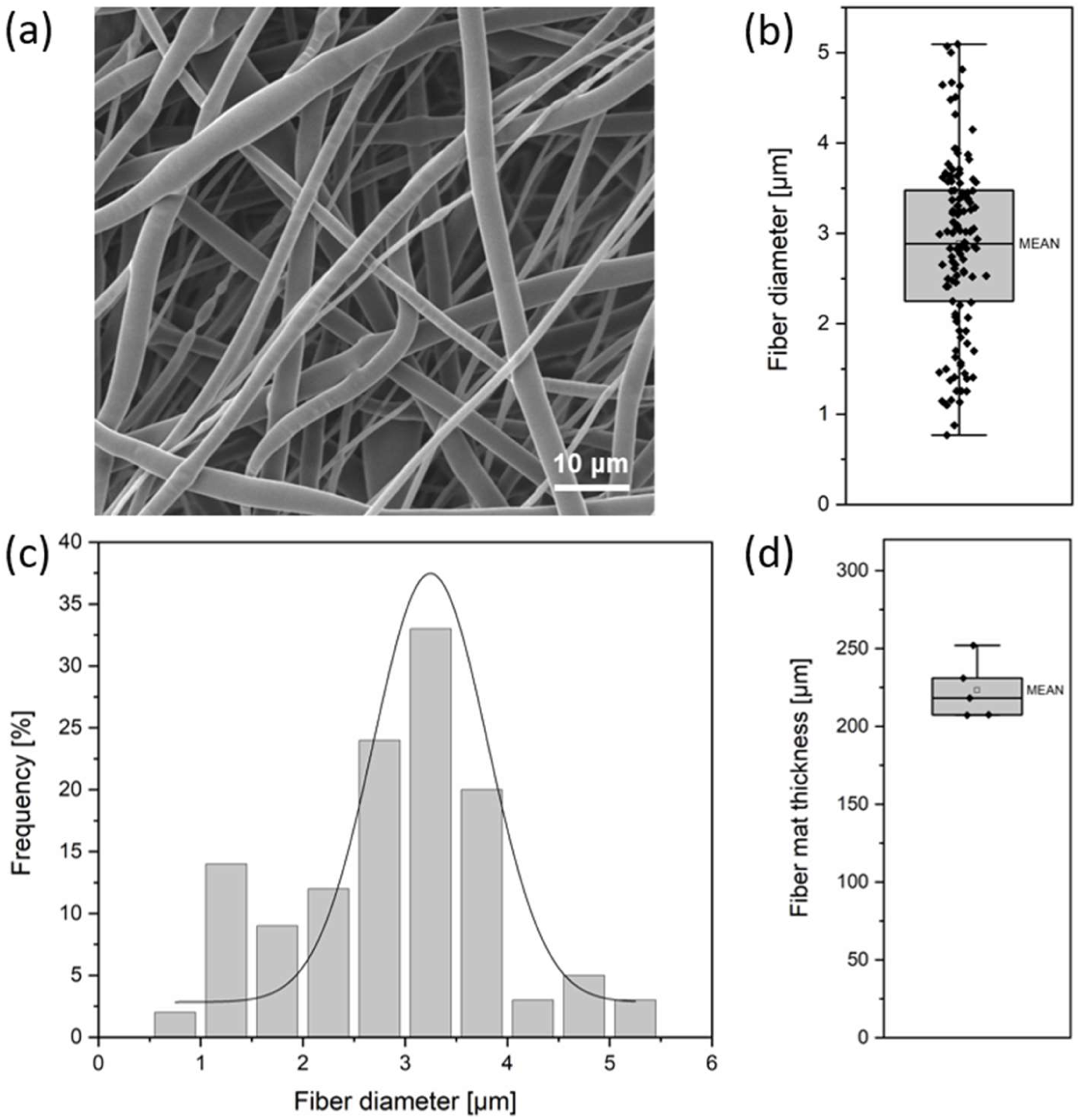
– Physical characterization of electrospun PCL scaffolds. (a) Visualization of fiber morphology using SEM imaging. (b) Absolute fiber diameter distribution in the low micrometer range. (c) Bar graph of percentage fiber diameter distribution with Gaussian fit. (d) Bar graph of the fiber math thickness.

### Combination Bioprinting and Electrospinning

For bioprinting a custom-built system based on a commercial consumer-level stereolithographic 3D printer was used. A custom printing platform constituted by an array of pillars fitting a 24-well plate was used (see **Figure 10**). Each pillar works as a bioprinting support in each individual well. Therefore, during a single run 24 individual constructs can be potentially printed (**Figure 2** and Materials and Methods section for additional technical details). To realize the hybrid system, a disk of PCL nanofiber was attached to a pillar. Next, the bioprinting was started. The hybrid constructs were composed of successive layers of 3D printed hydrogel attached to PCL nanofiber, with a total of 12 layers of printed hydrogel, forming discs with a diameter of 8mm and 2 mm in height (**Figure 2a**, **Figure 2b**). The attachment of these two types of scaffolds was stable in PBS at 37°C.

**Figure 2.**
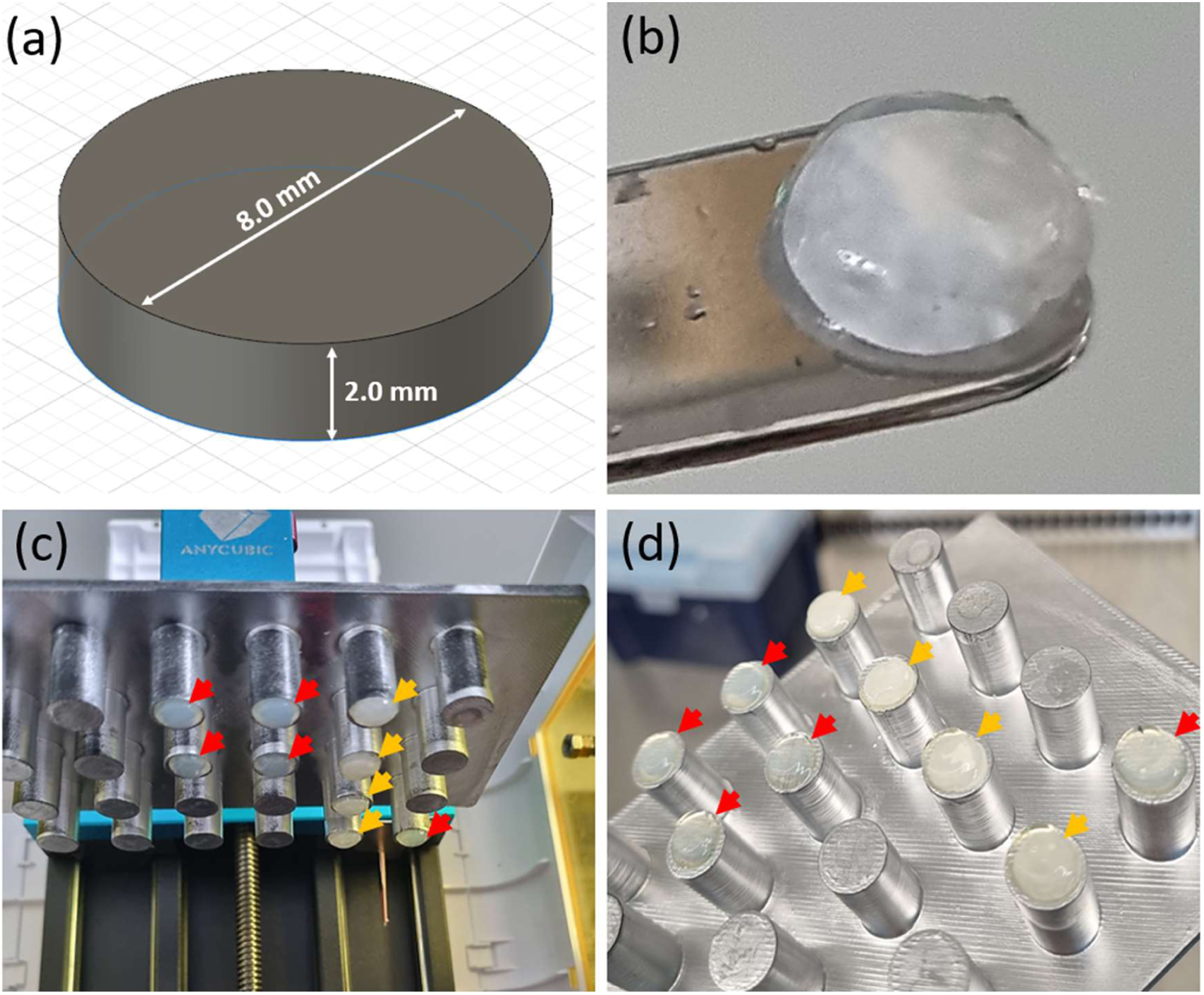
– Fabrication of hybrid construct (a) The CAD-designed of disc for bioprinting, measuring 8 mm in diameter and 2 mm in height (b) Hybrid construct composed of PCL nanofibers and a printed GelMA/PEGDA hydrogel disc (14%/6.1%, respectively) (c, d) Bioprinter platform after printing hydrogel on top of PCL electrospun mats. The orange arrows show hybrid constructs and the red arrows indicate printed hydrogel discs.

### Visualization of hydrogel/fiber hybrid constructs

Cross-sectional structure analysis of the hybrid constructs by SEM imaging revealed a clear discrimination between the two layers. More specifically, the transition from the smooth, porous hydrogel with interconnected pores to the dense, fibrous sheet on top could be visualized as represented in **Figure 3a**. In detail, the interfacial region crucial for the mechanical stability and functionality of the hybrid constructs being highlighted in the close-up SEM micrograph in **Figure 1a** demonstrated interfacial bonding of the two layers by interpenetrating of the hydrogel into the fibrous network. This led to embedding of the fiber top sheet into the upper hydrogel compartment creating a gradual transition zone rather than a sharp interface, indicating an interconnection of both compartments within the composite construct. From the “bottom perspective” of the hybrid construct, the hydrogel surface typically exhibited a homogeneous, smooth and continuous texture as shown in **Figure 3b**. Considering the top view side of the hybrid constructs with focus on the fiber scaffold, the nonwoven 3D fiber network appeared to be embedded into the hydrogel surface as depicted in **Figure 3c**.

**Figure 3.**
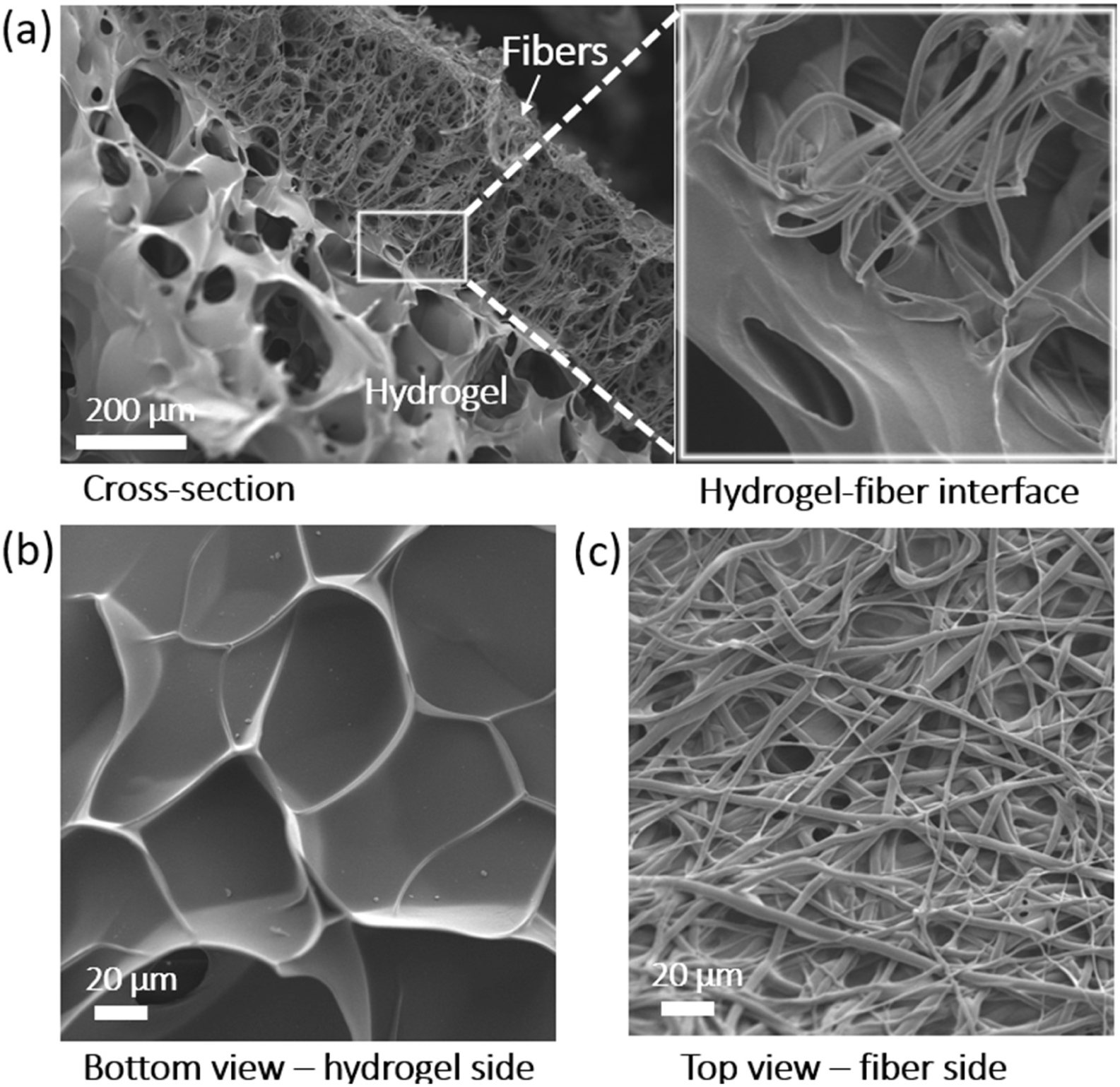
– Visualization of the morphology of the hydrogel/fiber hybrid constructs using SEM imaging. (a) Cross-sectional structure of hybrid constructs comprising a bioprinted hydrogel layer and an electrospun fiber top sheet highlighting the interconnected hydrogel-fiber interface as represented in the close-up view. (b) Bottom view representing the hydrogel side. (c) Top view representing the fiber side.

### Embedding of the hMSC spheroids in the bioprinted hydrogel

A large number of uniformly sized BM-hMSC spheroids was generated by using commercially available micro-well plates (**Figure 4a**). Although the spheroids were homogeneously mixed in the GelMA/PEGDA hydrogel bioink, during bioprinting they showed the tendency to accumulate at the periphery of the printed construct, likely due to capillary effects. We obtained a density of 30 to 50 spheroids per construct (**Figure 4b**).

**Figure 4.**
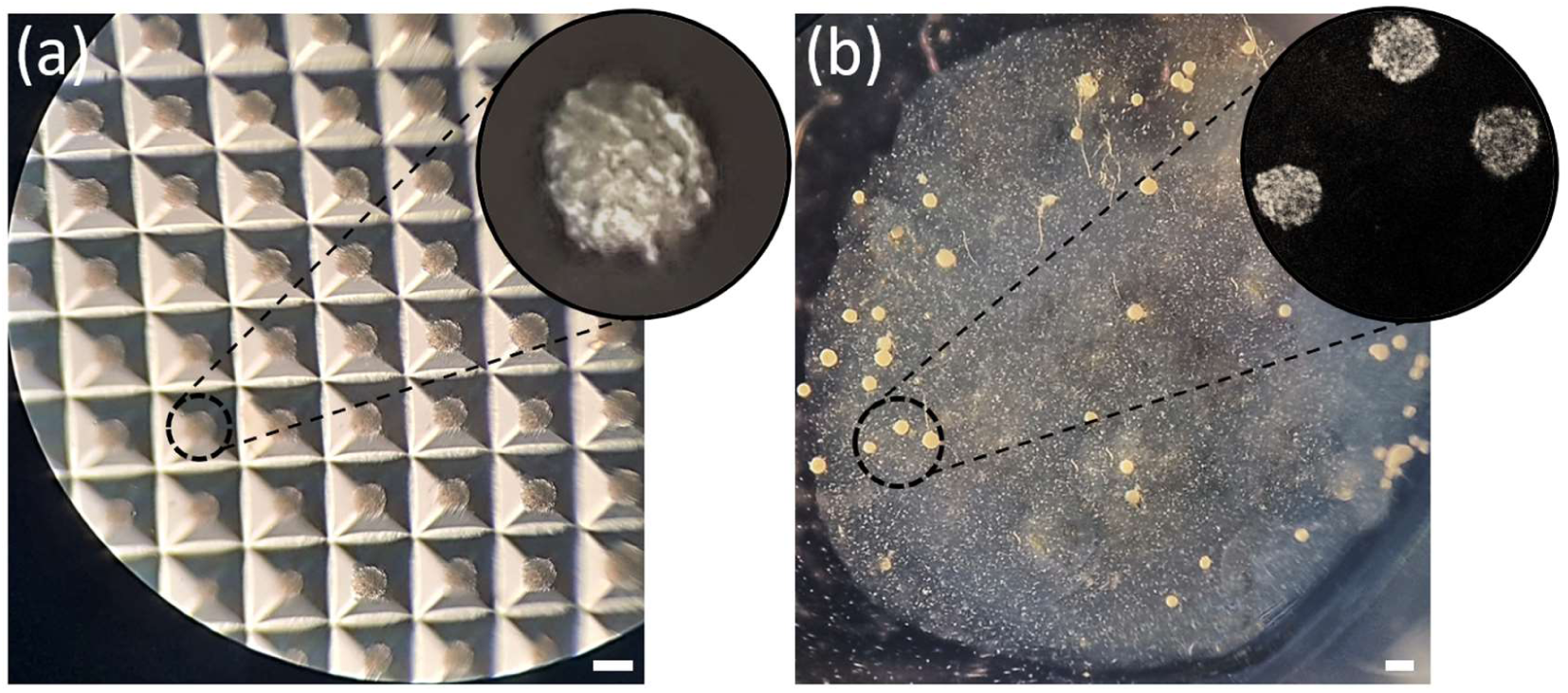
– (a) Formed spheroids in the 6-well Sphericalplate 5D after one day. Each one of the well contains 750 microwells, resulting in 750 spheroids per well, with 1,000 BM-hMSCs per microwell. The average size of the spheroids was measured at 130 µm by randomly selecting and averaging the values from 10-15 spheroids. (b) Printed spheroids in 14% GelMA/6.1% PEGDA hydrogel. 30-50 spheroids were embedded into an 8 mm by 2 mm printed disc. Scale bar: 200 µm. Microscope: Zeiss SteREO Discovery V8. Objective: Plan Apo S, 0.63×/0.116 FWD 81 mm. Camera: AxioCam IcC SIN. Pixel size: 4.54 × 4.54 µm².

### Live/dead assay

The viability of the BM-hMSC single cells vs. spheroids embedded in the bioprinted constructs was assessed on days 5 and 21 after bioprinting (**Figure 5**). On day 5, spheroids exhibited partial cell death within the central core, yet the majority of cells remained viable and spread throughout the construct. Remarkably, the sprouting cells proliferated across the entire construct and penetrated multiple layers (**Figure 6**). In comparison, isolated hMSCs displayed cell death and failed to attach to the printed hydrogel by day 5. However, they were able to spread by day 21, albeit limited to the surface of the construct (**Figure 5**). In contrast, the cells migrating from the spheroids thoroughly populated the entire construct (**Figure 6**).

**Figure 5.**
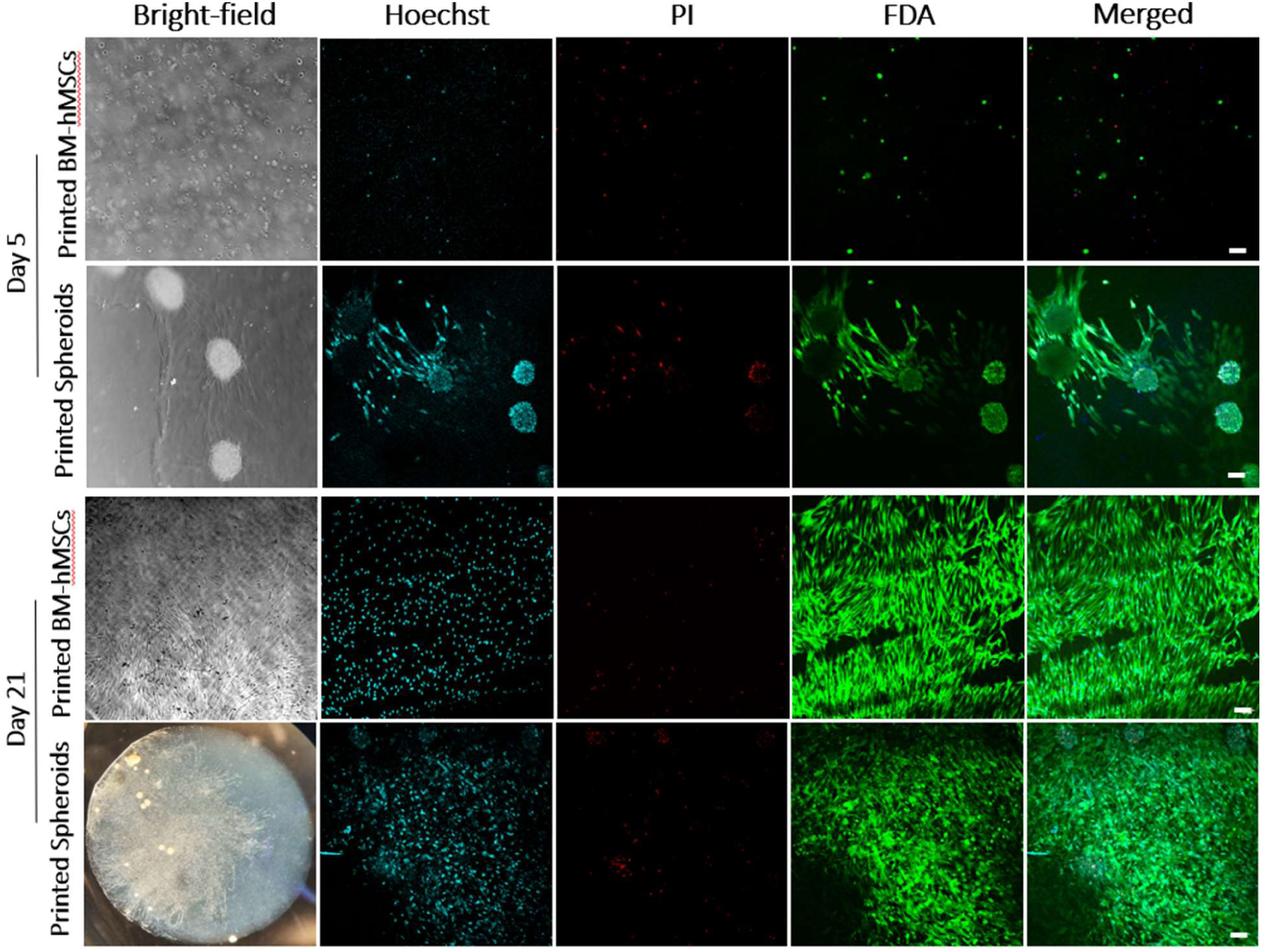
– Live-dead assay demonstrating the cell viability of bioprinted BM-hMSCs and spheroids at 5 and 21 days post-bioprinting. BM-hMSCs migrated out of the spheroids within the printed hydrogel. In contrast, single BM-hMSCs retained a round shape and did not develop the typical spindle-shaped morphology characteristic of BM-hMSCs. Live cells were stained with fluorescein diacetate (FDA, green channel), while dead cells were stained with propidium iodide (PI, red channel). Scale bar: 100 µm. Bright-field imaging was conducted using an inverted Zeiss Axiovert microscope, with additional imaging of the printed spheroids on day 21 performed using a Zeiss stereoscope.

**Figure 6.**
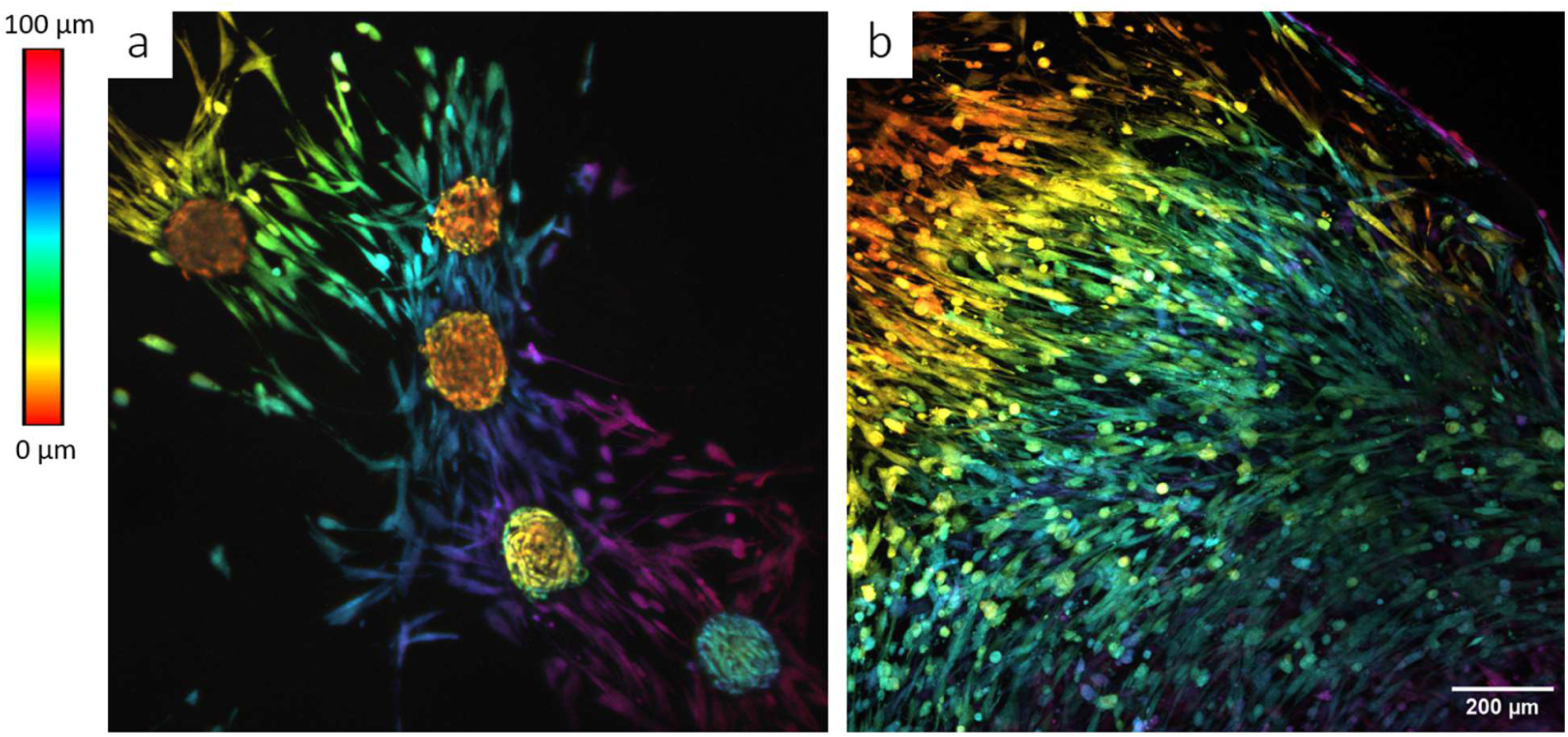
– (a) Bioprinted hMSC spheroids after 5 days of culture. Maximum projection of an image stack recorded with the confocal microscope. The color coding of the indicates the depth inside the construct in the range 0-100 µm. After 5 days, hMSCs are sprouting out of the spheroids and distributing across the whole construct depth. (b) appearance of the sample shown in (a) after 21 days of culture: hMSC cells evenly distributed inside the hydrogel at different depths. Images recorded with a Zeiss LSM780 confocal microscope, ZEISS Objektiv Plan-Apochromat 10x/0,3 M27. The depth color coding was performed with the ImageJ plugin “Z-Stack Depth Color Code” (https://github.com/UU-cellbiology/ZstackDepthColorCode).

### MTT assay on fibroblasts and keratinocytes

The MTT assay was conducted to evaluate the potential cytotoxic effects of the hydrogels on the viability of fibroblasts (Hs27) and keratinocytes (HaCaT) at both high– and low-glucose conditions after 24 hours. As shown in **Figure 7**, the viability assay results indicated no cytotoxic effects in the control, hydrogel only, and hybrid construct only groups. Conversely, cell viability in the printed BM-hMSC spheroid and BM-hMSC single cells groups significantly increased under low-glucose conditions (119.34% ± 8.33 and 112.86% ± 6.55 for Hs27; 130.71% ± 1.98 and 124.78% ± 14.58 for HaCaT, respectively). Additionally, printed spheroids notably enhanced the viability of fibroblasts under high glucose conditions (113.66% ± 3.47).

**Figure 7.**
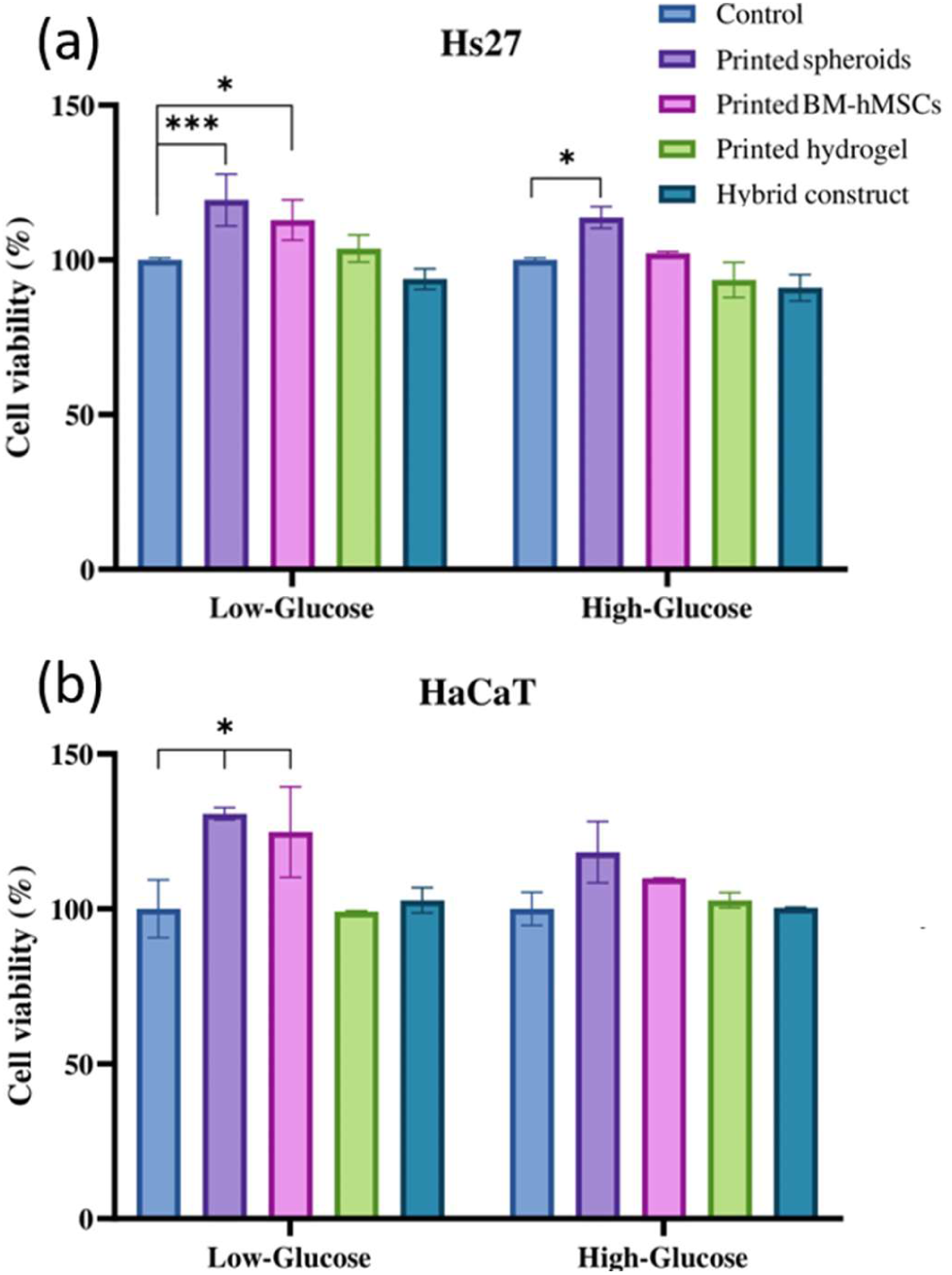
– MTT assay, the relative cell viability of constructs on (a) human fibroblasts (Hs27) and (b) keratinocytes (HaCaT) under normal and high glucose conditions at 24h post-incubation. Data are expressed as mean ± SD, n = 3 (*p < 0.05, and ***p < 0.001).

**Figure 7.**
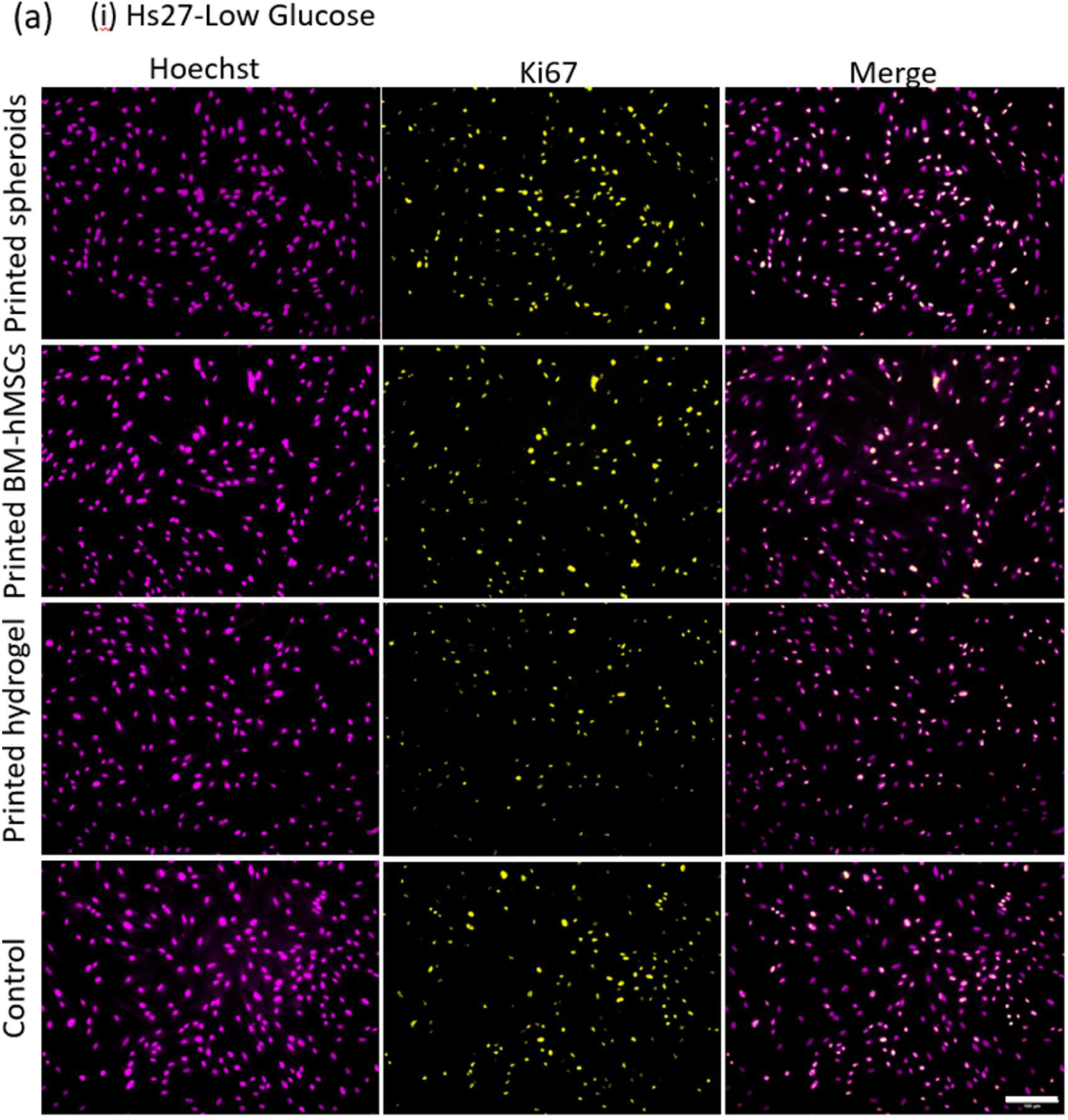
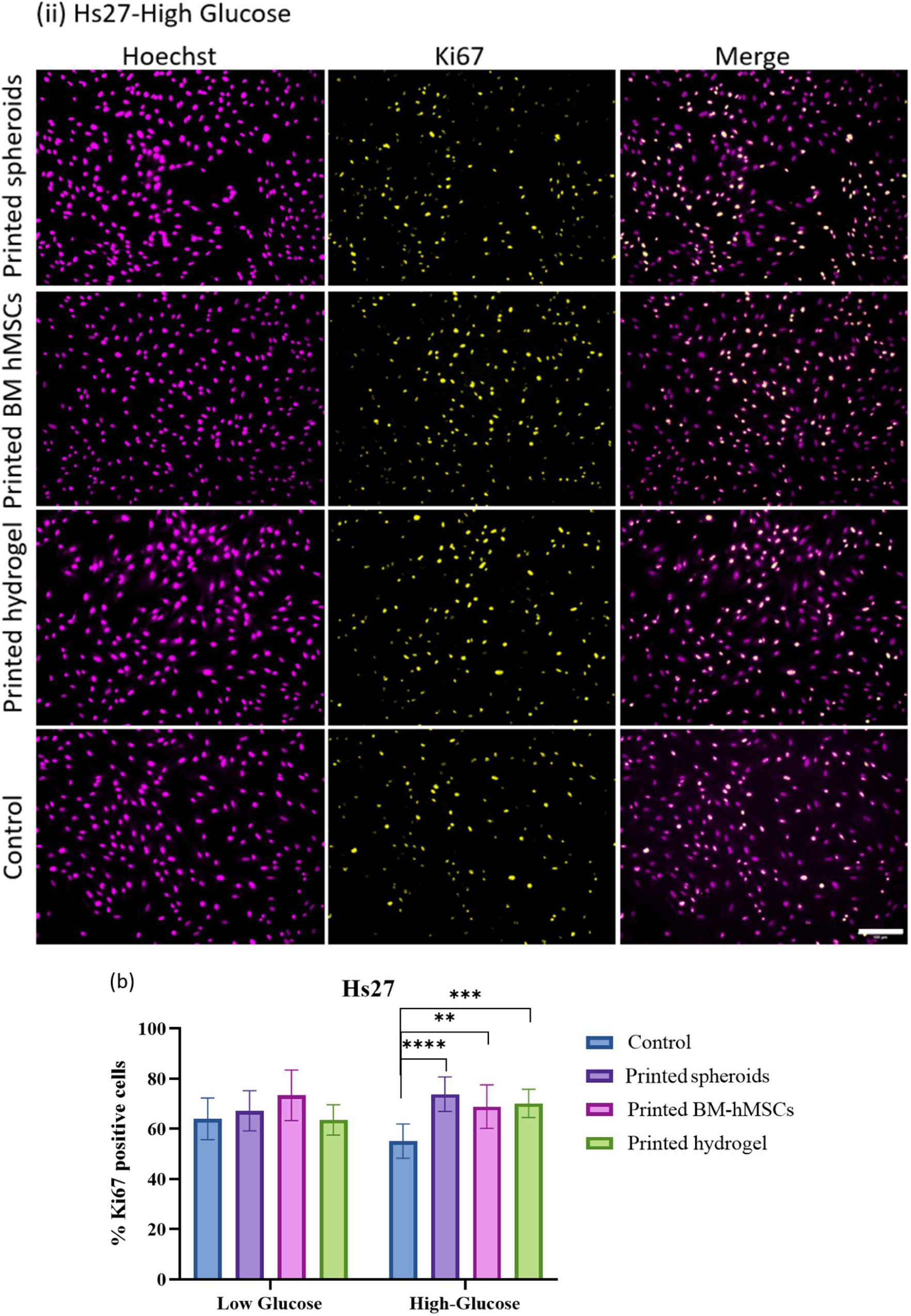
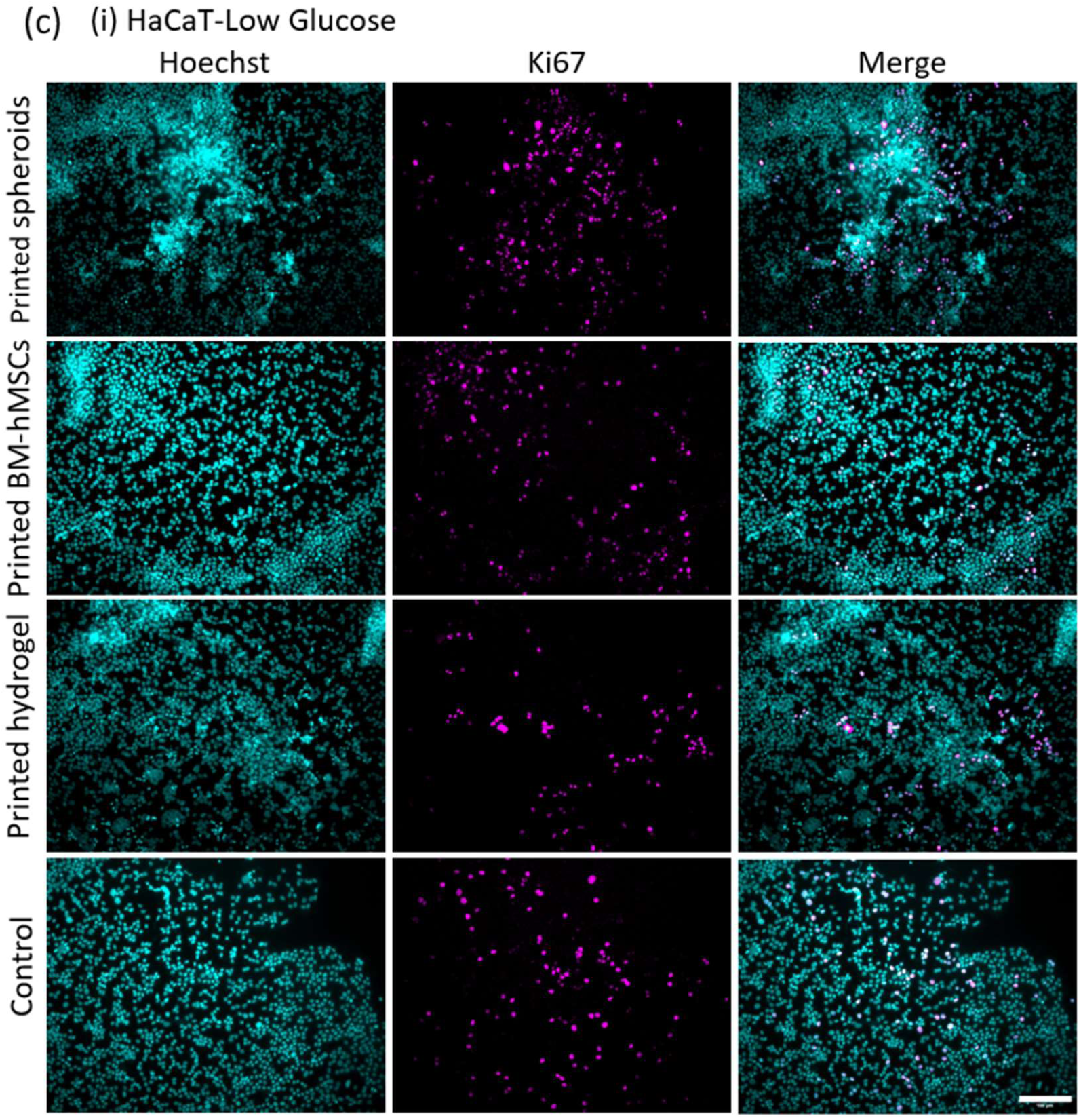
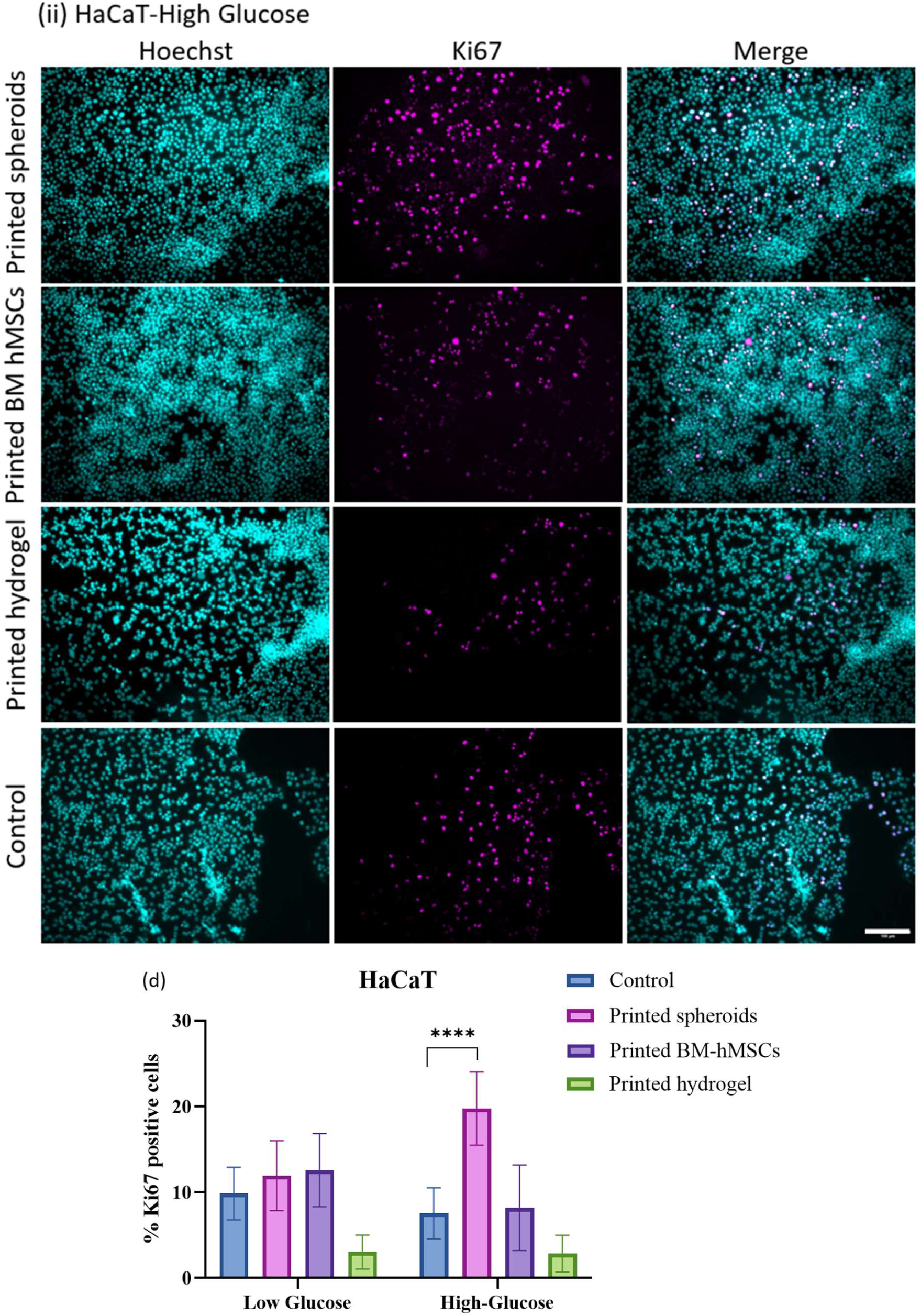
– Presence of the proliferation marker Ki67 in (a) human fibroblasts Hs27 and (c) HaCaT keratinocytes incubated with printed constructs under both (i) low and (ii) high glucose conditions. (b)The quantification of Ki67-positive shows that all treatment groups exhibited a proliferative effect on human fibroblasts under high glucose conditions. (d) A higher number of Ki67-positive keratinocytes was observed under high glucose conditions in the printed spheroid group. Images of representative constructs stained for Ki67 (yellow, λ_ex: 488 nm, λ_em: 490–562 nm) and Hoechst 33342 for nuclei (magenta, λ_ex: 405 nm, λ_em: 410–502 nm) are shown. (b), (d) Quantification of the ratio of proliferating cells (Ki67-positive) to the total number of cells (Hoechst 33342-stained). Data are presented as mean ± SD, n = 7 (**p < 0.01, ***p < 0.001, ****p < 0.001).

### Cell proliferation

Immunostaining with the Ki67 antibody, a well-established marker for cell proliferation, was used to assess the proliferative response in various treatment groups. In the low-glucose condition, BM-hMSCs and spheroids within the bioprinted constructs promoted cell proliferation in both fibroblast and keratinocyte cell lines compared to controls (**Figure 7b**, **Figure 7d**). However, a slight reduction in the ratio of proliferating cells to the total cell count was observed in the printed hydrogel group under these conditions. On the other hand, in the high-glucose conditions, a significant increase in fibroblast proliferation was observed in the treatment groups (**Figure 7b**). Additionally, the percentage of Ki67-positive keratinocytes was markedly elevated in groups exposed to bioprinted BM-hMSC spheroids under these conditions (**Figure 7d**).”

### Scratch assay

To mimic wound healing *in vitro* and assess whether the constructs promote cell migration during wound closure, confluent HaCaT and Hs27 cells were incubated for 36-48 hours. The wound margin was marked, and the migration rate was quantified every 12 hours of incubation. The results demonstrated that both HaCaT and Hs27 cells exhibited higher migration rates in the presence of printed spheroids compared to the medium alone. At 36 hours, the closure in Hs27 scratch assays was 83.73 ± 12.15% under low glucose and 98.4 ± 1.93% under high glucose, and 80.95 ± 17.14% under low glucose and 66.44 ± 7.83% under high glucose in the printed BM-hMSCs group. The control group showed wound closure rates of 56.7 ± 11.5% under low glucose and 58.42 ± 9.27% under high glucose (**Figure 8a, b**). Therefore, the scratch wound assay demonstrated that spheroids and BM-hMSCs could significantly increase the rate of cell wound closure compared with a medium in vitro under low glucose level (p < 0.01), and only spheroids could enhance cell migration under high glucose (****p < 0.001).

**Figure 8.**
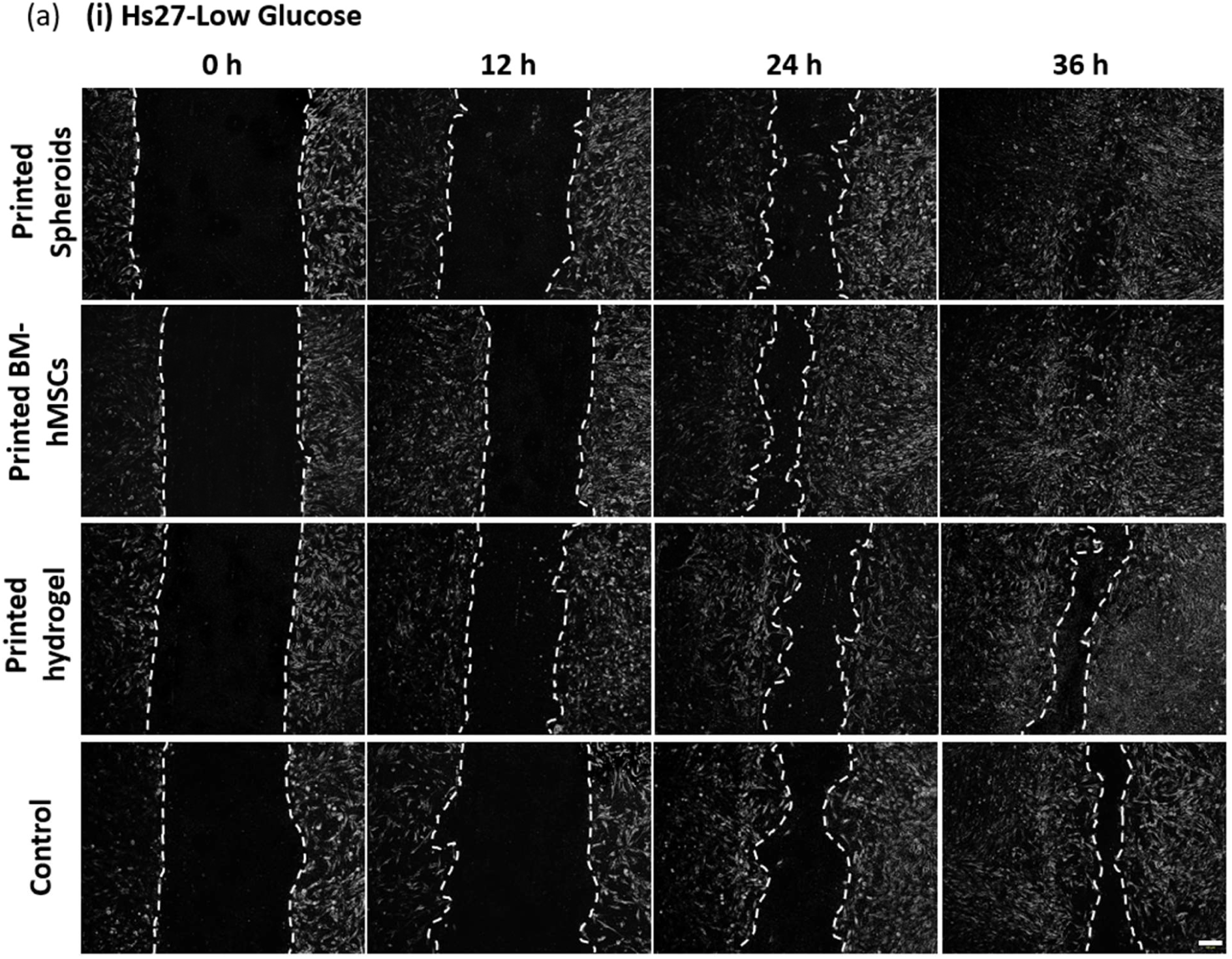
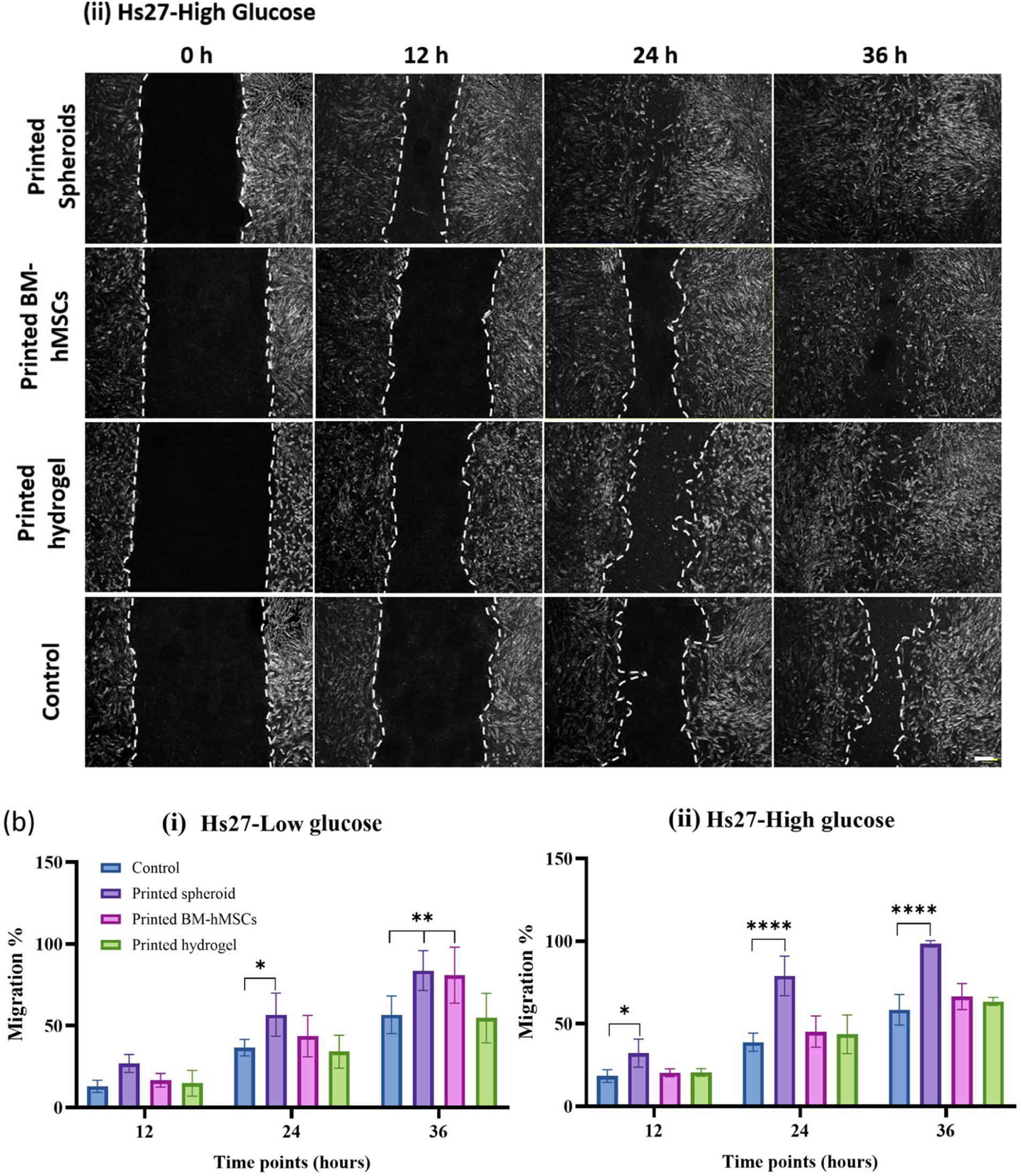
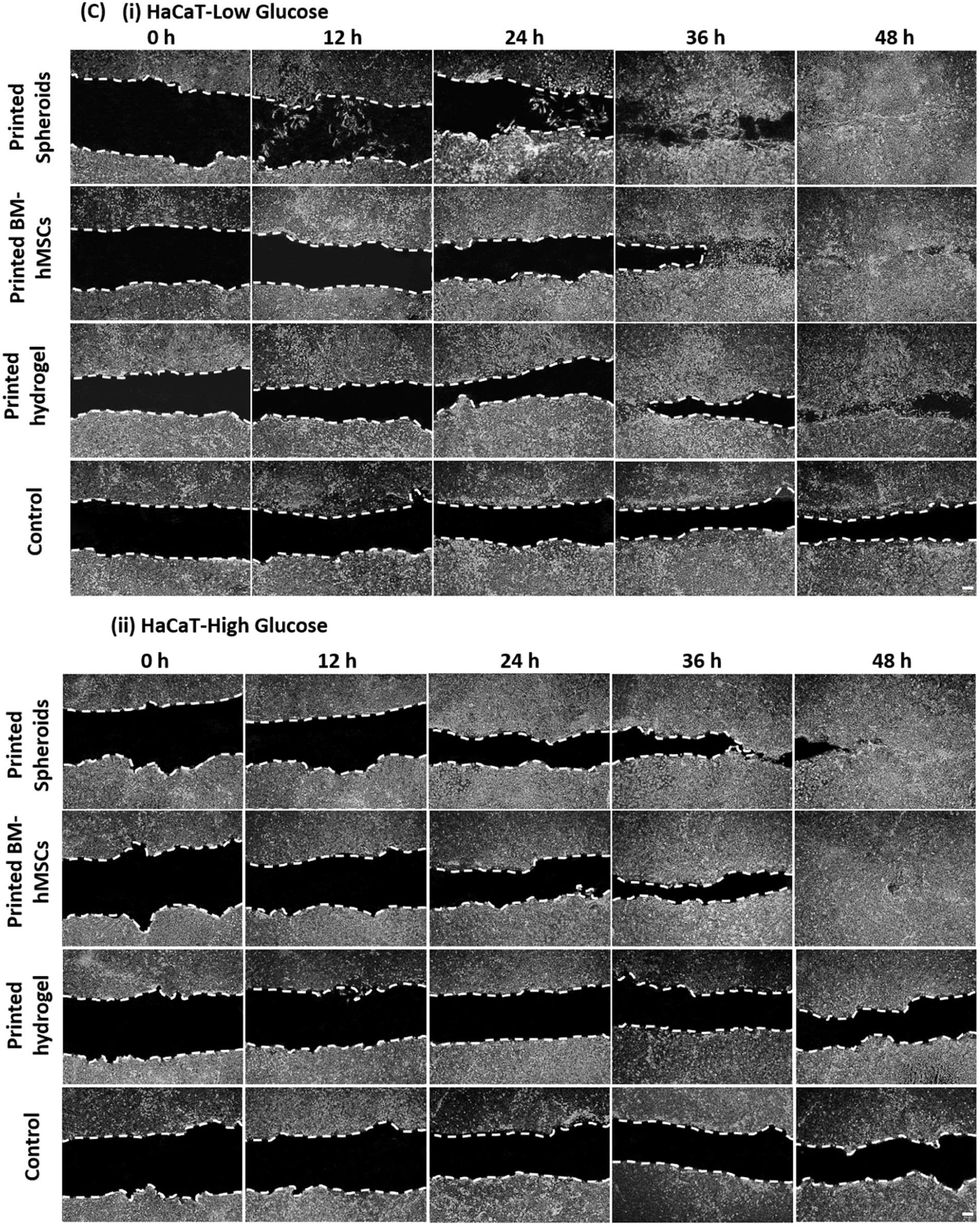
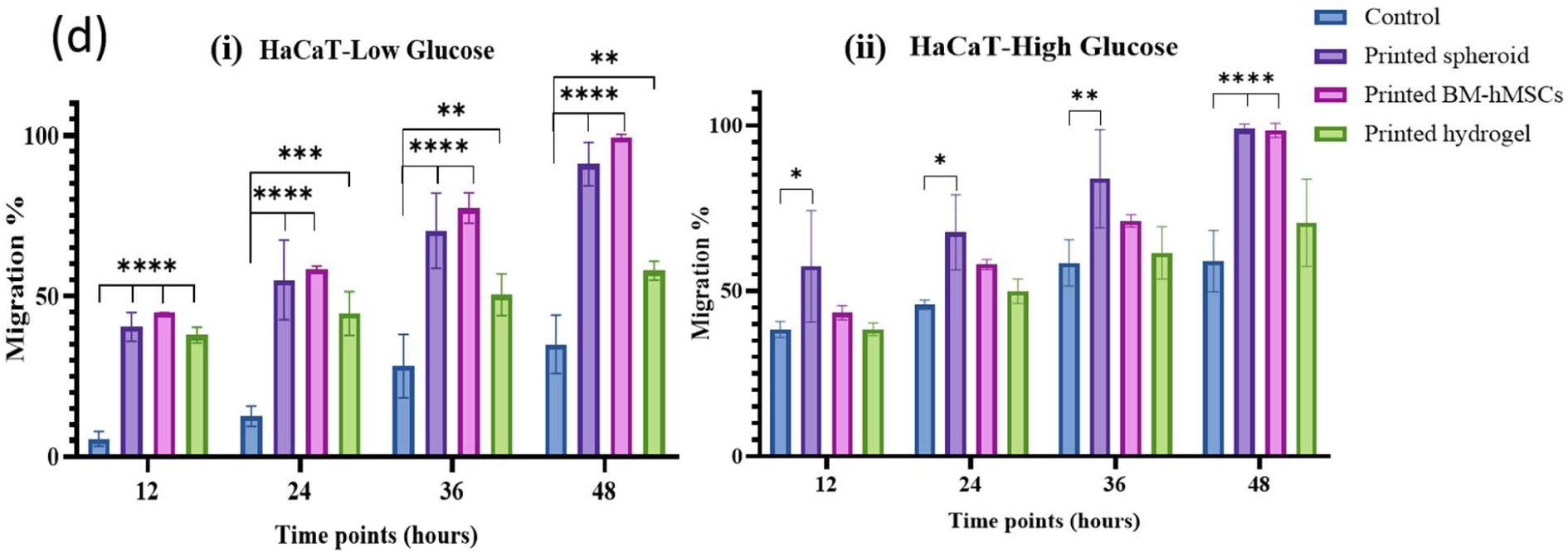
– (a) The microscopic images of the untreated human fibroblasts (Hs27) and treated with the printed spheroids, printed BM-hMSCs, and printed hydrogel after 12h, 24 h, and 36h under (i) low and (ii) high glucose levels. (b) The microscopic images of the untreated human keratinocyte (HaCaT) and treated with the printed spheroids, printed BM-hMSCs, and printed hydrogel after 12h, 24 h, 36h, and 48h under (i) low and (ii) high glucose levels. In each image, the leading edges are plotted in yellow. Scale bar: 200 μm. (b,c, and d) The quantitative assessment of the percentage of cell mobility after the incubation with or without the constructs. Data are shown as mean ± SD, n = 3 (*p < 0.05 **p < 0.01, ***p < 0.001, ****p < 0.001).

Similarly, for HaCaT cells at 48 hours, wound closure in the printed spheroids group was 91.05 ± 6.78% under low glucose and 99 ± 1.41% under high glucose, and in the printed BM-hMSCs group 99.03 ± 0.97% under low glucose and 98.5 ± 2.12% under high glucose. The control group showed wound closure rates of 56.7 ± 11.5% under low glucose and 58.42 ± 9.27% under high glucose (**Figure 8c, d**). A significant increase was present for both groups and conditions compared to control (****p < 0.001).

On the other hand, the findings of the HaCaT scratch assay (detectable due to the different morphology of HaCaT and BM-hMSCs) indicated that BM-hMSCs could migrate from the printed spheroid construct to the wound scratch site and support wound closure (Figure S?, Supporting Information).

## Discussion and outlook

In this study, we successfully developed a hybrid construct combining two advanced fabrication techniques—electrospinning and light-based bioprinting—and two distinct materials, polycaprolactone (PCL) for mechanical strength and a hydrogel blend of polyethylene glycol methacrylate (PEGMA) and gelatin methacrylate (GelMA) for enhanced biocompatibility and cellular proliferation. The construct was designed to address the persistent challenge of diabetic wound healing, offering a functional scaffold capable of promoting tissue regeneration. The results presented in this study demonstrate the potential of our hybrid construct in several key areas: mechanical properties, cell viability, and wound closure efficacy. The inclusion of PCL nanofibers significantly enhanced the mechanical stability of the scaffold, facilitating handling and deposition on the model wound. This addresses a common limitation of pure hydrogel-based constructs, which are difficult to manipulate due to their low Young’s modulus. The stability of the PCL and hydrogel attachment is crucial for the scaffold’s performance under physiological conditions, ensuring it can withstand mechanical stresses during implantation and use. Future studies should aim to quantify the mechanical properties more precisely through tensile and compressive testing to provide a comprehensive understanding of the construct’s behavior under different stress conditions.

The use of mesenchymal stem cell (MSC) spheroids as cellular reservoirs within the hydrogel matrix proved advantageous for cell viability and proliferation. The spheroids maintained a uniform spherical morphology and due to rapid settling during bioprinting they were predominantly distributed near the periphery of the construct, with a density of 30 to 50 spheroids per construct. The live/dead assay revealed that while there was partial cell death within the core of the spheroids on day 5, the majority of cells remained viable and spread throughout the construct by day 21. This placement facilitates efficient nutrient diffusion and waste removal, contributing to enhanced cell survival and proliferation during the construct’s maturation. Our findings underscore the importance of spheroid formation in promoting cell viability over extended culture and maturation periods. In comparison, isolated MSCs exhibited higher cell death rates and limited attachment in the construct, highlighting the superiority of the spheroid approach. Nevertheless, optimizing spheroid size and density could further improve cell viability and distribution within the scaffold.

The scratch assay results demonstrated that both human fibroblasts (Hs27) and keratinocytes (HaCaT) exhibited higher migration rates in the presence of the printed spheroids construct compared to controls. The enhanced wound closure observed in both low– and high-glucose conditions suggests that the hybrid construct can effectively support cell migration and tissue regeneration in diabetic environments. Specifically, the significant increase in wound closure rates (up to 98.4% for fibroblasts and 99% for keratinocytes) under high glucose conditions is particularly promising, given the impaired healing associated with diabetes.

The MTT assay results confirmed that the hydrogel components were non-cytotoxic, with printed spheroids and BM-hMSCs significantly enhancing cell viability under normal and high-glucose conditions. This biocompatibility is critical for clinical applications, ensuring that the scaffold does not elicit adverse immune responses upon implantation.

While the current study shows the potential of our approach combining hybrid electrospun, and bioprinted scaffolds, and cellular spheroid seeding in wound healing, several areas require further investigation. Conducting long-term *in vivo* studies will be crucial to assess the scaffold’s performance in a more complex biological environment. These studies should evaluate not only wound healing efficacy but also potential immune responses and biodegradation rates. Future work should focus on optimizing the scaffold design, including variations in layer composition, thickness, and porosity. These parameters could be fine-tuned to enhance nutrient diffusion, cell migration, and mechanical properties. Exploring the integration of this scaffold with other therapeutic modalities, such as growth factors or antimicrobial agents, could further enhance its efficacy in treating chronic wounds. A further interesting development would involve varying the shape of the bioprinted hydrogel to test how different structures influence the performance of this wound dressing. For instance, incorporating triangular or parallelepipedal motifs could improve efficacy by increasing the contact surface area (**Figure 9**). These investigations will be conducted in follow-up studies.

**Figure 9.**
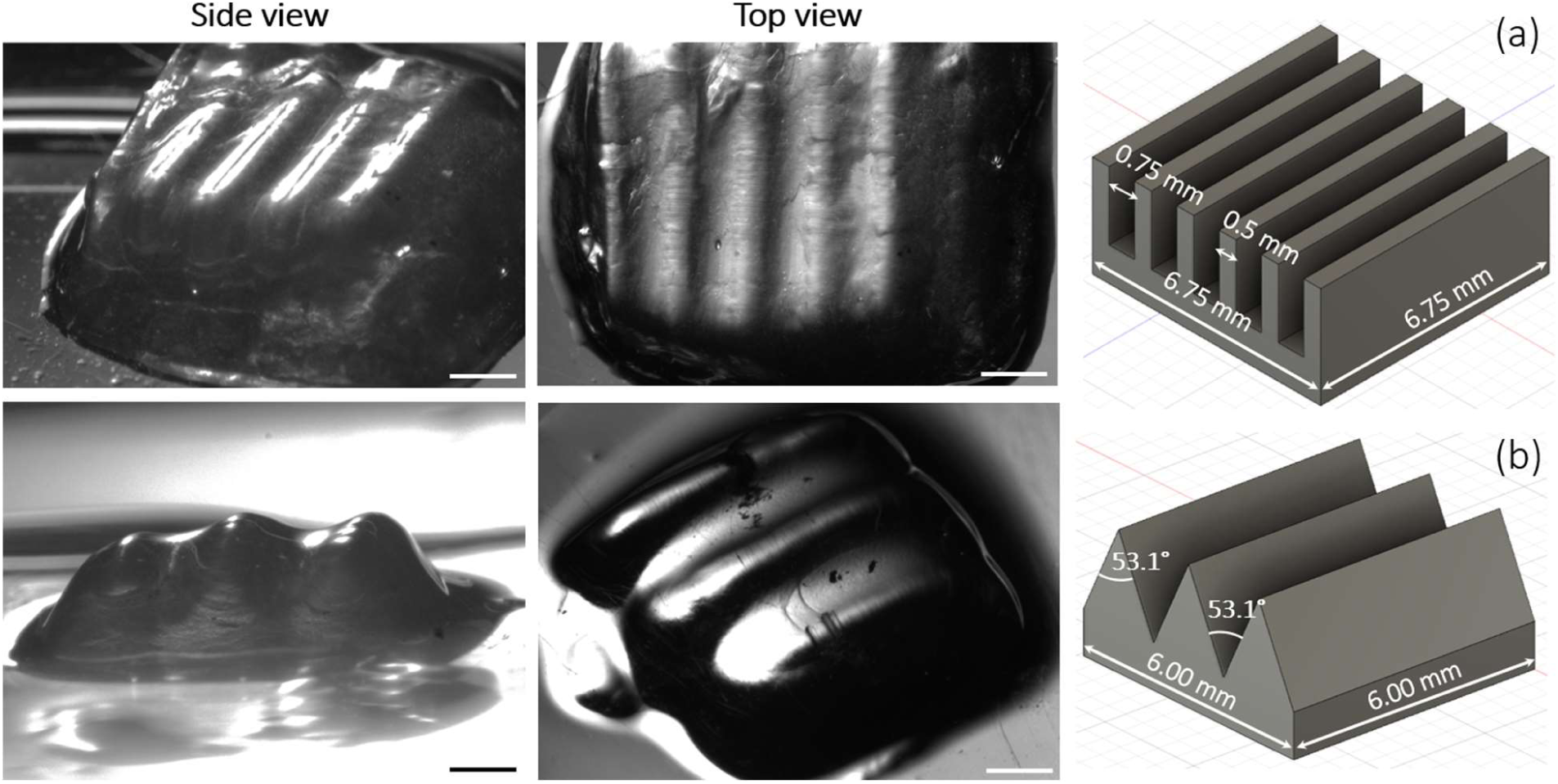
– 3D bioprinted constructs of variable shapes (square and triangle) for different applications using 7%/3% GelMA/PEGDA hydrogel. Right column: CAD design, middle and left column: stereomicroscope images. A) Square shape and B) triangle shape. Scale bar: 1mm. Microscope: Zeiss SteREO Discovery V8. Objective: Plan Apo S, 0.63×/0.116 FWD 81 mm. Camera: AxioCam IcC SIN. Pixel size: 4.54 × 4.54 μm^2^.

In conclusion, the hybrid construct developed in this study represents a promising step forward in realizing bioengineered wound dressing. By combining electrospinning and bioprinting techniques, we have created a scaffold that addresses both mechanical and biological, requirements for effective wound healing.

## Materials and Methods

### Mesenchymal stem cell culture and spheroid formation

BM-hMSCs were purchased from the company PromoCell (Heidelberg, Germany) and cultured in the growth medium at 37 °C in an atmosphere of 5% CO^2^. BM-hMSCs with 80% confluency between passages 3 and 7 were used for subsequent experiments.

Spheroid formation: the BM-hMSC spheroids were formed in “Sphericalplate 5D” multi-well plates (Kugelmeiers, Erlenbach, Switzerland) following the protocol provided by the manufacturer. Briefly, 1 ml culture media was added to each well of Sphericalplate 5D plate to make sure there was no bubble. Then, 750000 BM-hMSCs (1000 cells per microwell) were carefully added to each well. The seeded cells were transferred to the cell culture incubator. The condition media was exchanged every other day for 4 days.

### Preparation of photo-crosslinkable hydrogels and cell/spheroid encapsulation

Hydrogel preparation was carried out according to a previous study (16). The hydrogel used for this study was a combination of gelatin methacrylate (GelMA) gel strength 300 g, Bloom, 80% degree of substitution (Sigma Aldrich Chemie GmbH, Steinheim, Germany), and polyethylene glycol diacrylate (PEGDA), average Mn 4000 (Sigma Aldrich Chemie GmbH, Steinheim, Germany). The hydrogel components were separately mixed with either phosphate buffer saline (PBS, Gibco, ThermoFisher Scientific, Waltham, MA, USA). 0.2% w/v Lithium-Phenyl-2,4,6-trimethylbenzoylphosphinat (LAP, Sigma Aldrich Chemie GmbH, Steinheim, Germany) was added to the PEGDA mixture and the GelMA and PEGDA/LAP mixtures were dissolved separately at 65 °C under shaking conditions for 2 h. Subsequently, the dissolved GelMA and PEGDA were mixed to reach the desired concentration, 0.0025% w/v tartrazine (Sigma Aldrich Chemie GmbH, Steinheim, Germany) was added, and the mixtures were left at 37 °C for another hour. The concentrations used in this study were 14% w/v GelMA, 6.1% w/v PEGDA, and 0. 41% LAP. To encapsulate the cells in the hydrogel before 3D bioprinting, the BM-hMSCs were detached from the flask using Accutase and collected by centrifugation in a pellet (220 rcf, 3 minutes). The supernatant was discarded, and the cells were resuspended in the previously prepared hydrogel (see previous sections) with a density of 333000 cells/ml. To encapsulate spheroids a density of 325 spheroids/ml was applied. The sample/hydrogel mixtures were pipetted into the 24-well plate. After bioprinting, the construct was extracted using a metal spatula. The bioprinted objects were washed in PBS supplemented with 2% pen/strep to prevent potential contamination linked to handling and were subsequently cultured in a well plate for 24 days. The BM-hMSCs and spheroids within the printed hydrogel on day 14 were utilized for cell assays.

### Bioprinting design

Bioprinting is conducted by using a custom-built stereolithographic bioprinter (SLA bioprinter) (16) that allows the layer-by-layer polymerization of a photo-crosslinkable GelMA-PEGDA hydrogel. To be able to bioprint in a format compatible with multi-well plates, a special printing platform suitable for 24-well plates was designed and 3D printed with a Bambu Lab X1 filament 3D printer using a PLA filament. The PLA platform was sprayed with aluminum to improve the adherence of the hydrogel on the platform during the printing process (**Figure 10**, **Figure 11**).

**Figure 10.**
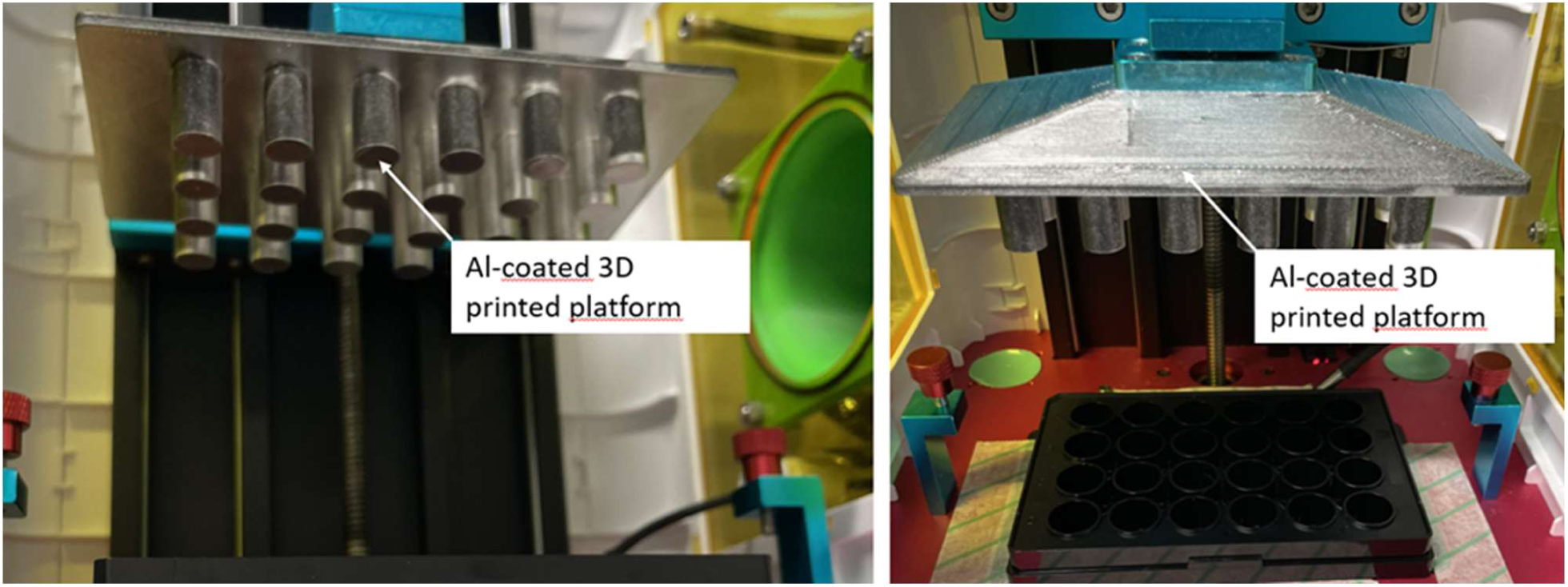
– Custom 3D printing plate compatible with a 24-well format. The plate is shown as already installed in the bioprinter.

**Figure 11.**
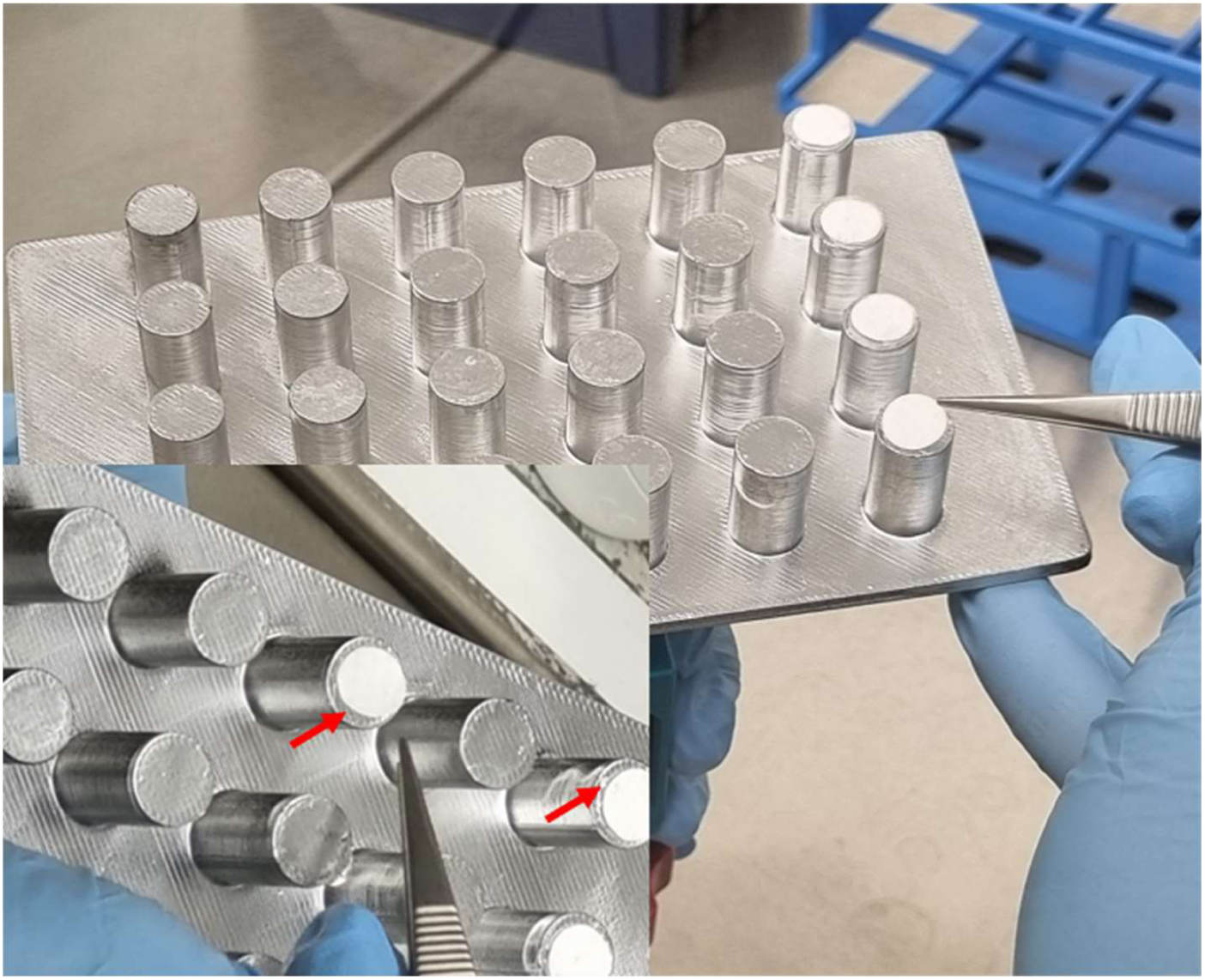
– Attachment of the electrospun PCL scaffold on the bioprinting platforms. In order to ensure a stable yet biocompatible adhesion of the scaffold with the platform, a droplet of agarose is used as a “glue” between both materials. After gelation, the agarose effectively keeps in place the PCL substrate during the bioprinting.

To print the Hydrogel Samples, the constructs were all designed with Fusion 360 (Autodesk Inc., San Rafael, CA, USA) computer-aided design (CAD) software and sliced with Chitubox (Chitubox, Shenzhen, Guangdong, China). The constructs encapsulating BM-hMSCs and BM-hMSC spheroids were disc-shaped 2 mm in height, and 8 mm in diameter. The settings of the slicing software for printing were 150 s bottom illumination (3 layers), 100 s exposure time, 150 μm layer height, 3 mm lifting distance, and 9 mm s−1 speed (bottom lift, lift, and retract speed). Indeed, a slower stage lifting speed was shown to distribute the spheroids fragments more homogenously within the construct.

### Fabrication of electrospun fiber scaffolds

For the fabrication of electrospun fiber scaffolds, a polymer solution of 15 % (w/v) polycaprolactone (PCL, Mw 80 kDa, Sigma Aldrich, Darmstadt, Germany) dissolved in a 50:50 (v/v) mixture of chloroform and ethanol (pharma grade reagents from VWR Chemicals, Darmstadt, Germany) was prepared by stirring at room temperature overnight. Afterwards, the degassed polymer solution was transferred into a 10 ml syringe (B. Braun Melsungen AG, Melsungen, Germany) equipped with a 21-gauge blunt tip needle (B. Braun Melsungen AG, Melsungen, Germany). The electrospinning process was performed based on a custom-built horizontal cabinet setup. The prepared polymer solution was pumped through a syringe nozzle at a flow rate of 2.5 ml/h with an applied voltage of 13 kV under ambient temperature. The formed fibers were deposited on a metal drum collector rotating with 150 revolutions per minute (rpm). The fabricated fiber scaffolds were stored in a desiccator at room temperature until further use.

### Physical characterization of electrospun PCL nanofiber scaffolds

The thickness of the electrospun fiber scaffolds was determined on sample punches measuring 10 mm in diameter using the MiniTest 735 coating thickness gauge with a F5 sensor (ElektroPhysik Dr. Steingroever GmbH & Co. KG, Cologne, Germany). The scaffold punches were placed on a stainless-steel metal plate and covered with a previously tared microscopy cover slip to allow for uniform pressure distribution by placing the sensor on top, thus enabling automatic reading by magnetic induction. Five samples per batch were analyzed in fivefold technical determination.

### Printing bioink on the surface of electrospun PCL

To directly print bioink on PCL nanofibers, a thin layer of liquid agarose (1.5% w/v) was coated on the platform, and the PCL mat was immediately applied on top (**Figure 11**). After the agarose was gelled, the coated platform was installed in a 3D bioprinter, and the printing of the bioink was started.

### Visualization of electrospun PCL scaffolds and bioprinted hydrogel/fiber hybrid constructs

Visualization of the PCL fiber morphology and the structural appearance of the bioprinted hydrogel/fiber hybrid constructs was performed using an EVO 10 scanning electron microscopy (SEM) by Zeiss (Carl Zeiss Microscopy GmbH, Oberkochen, Germany) at an acceleration voltage of 7 kV. For SEM image acquisition, the sample punches were placed onto specimen stubs covered with a conductive carbon adhesive tape followed by gold / palladium surface sputter coating for 150 s (SC7620 Mini Sputter Coater, Quantum Design GmbH, Darmstadt, Germany). Average fiber diameter was assessed on SEM micrographs at 500x magnification by randomly selecting five regions of interest from each scaffold batch and measuring 25 individual fibers from each area (in total = 125 fibers) using Fiji software (National Institutes of Health, USA). Prior to SEM investigation, the hybrid constructs were frozen for 16 h at –80°C and in a non-dehydrated state being freeze-dried for approx. 58h (Epsilon 2-4 LSCplus, Martin Christ Gefriertrocknungsanlagen GmbH, Osterode am Harz, Germany).

### Live/dead assay

Dead/live assay: The viability of BM-hMSCs after bioprinting was assessed using propidium iodide (PI) and fluorescein diacetate (FDA) staining. The bioprinted constructs were incubated in BM-hMSC growth medium for 4 days, washed with warmed PBS, then incubated at 37°C for 15 minutes in medium without supplements and phenol red, that contained 1:100 PI (1mg/ml), 1:500 FDA (5mg/ml) and 1:500 Hoechst (nuclear stain). After incubation, the constructs were once more washed in PBS and imaged in the medium.

### High-glucose model

Human fibroblasts (Hs27) and human keratinocytes (HaCaT) were cultured at 37 °C with 95% humidity and 5% CO2 in Dulbecco’s Modified Eagle’s Medium (DMEM). For euglycemic conditions, both cell lines were maintained in DMEM containing 5.5 mM glucose. For hyperglycemic conditions, keratinocytes and fibroblasts were cultured in DMEM containing 25 mM and 50 mM glucose, respectively, for 48 hours prior to start the experiments.

### MTT assay

Microculture tetrazolium (MTT) assay was performed as elsewhere described to study the viability of fibroblasts and keratinocytes after incubation with constructs. Briefly, HaCaT and Hs27 (1×10^4^ and 1.4×10^4^ cells/cm^2^, respectively) were seeded into 24-well plates in triplicates and analysis was performed at a 24-hour time point. Thiazolyl Blue tetrazolium bromide (Sigma-Aldrich) was used at 0.5 mg/mL in culture media and the formazan crystals generated were dissolved in DMSO. Optical density was measured at 570 nm after subtraction of 650 nm background on a TECAN microplate reader (TECAN, Männedorf, Switzerland).

### Proliferation assay

Proliferating cells were stained against Ki67 and quantified by fluorescence microscopy. Briefly, cells were fixed in 4% PFA (MilliporeSigma, St. Louis, MO, United States) in PBS for 30 min. They were then washed three times in PBS and permeabilized in Triton X-100 (0.3% v/v) in PBS for 40 min at room temperature. After washing thrice with 0.1 m glycine (Carl Roth GmbH, Karlruhe, Germany) in PBS and thrice in PBS-T (0.1% Triton X-100 in PBS, MilliporeSigma, St. Louis, MO, United States) for 10 min each time, the samples were blocked for 1 h in BSA (0.1%), Triton x-100 (0.2%), Tween-20 (0.05%) in PBS and incubated in the primary antibody solution overnight (see Figure S10, Supporting Information). The next day, the samples were washed in 2% penicillin/streptomycin in PBS thrice for 5 min before incubating in the secondary antibody solution for 2 h. After washing, the nuclei were stained by Hoechst (1:500), and the cells were ready for imaging following a final wash in PBS and 2% penicillin/streptomycin. Two biological replicates were imaged per condition, with at least 3 images per biological replicate.

### Scratch assay

The migratory effect of constructs was evaluated by wound scratch assay on both Hs27 and HaCaT cell lines. Hs27 and HaCaT cells (50×10^3^ and 25×10^3^, respectively) were seeded in 24-well plates with low serum media (2% FBS). The cells were scratched with a pipette tip 1ml when reaching 90% confluence. A microscope was used to photograph cells at 0 h, 12 h, 24 h, and 36 h after the scratch. The distances between the two borders of the scratch were calculated through ImageJ software. The migration rate was calculated with the following formula: cell migration rate = (1 – blank space width (each time point) / blank space width (0 h)) ×100%.

## Acknowledgements

S.A.H. acknowledges funding by Mashhad University of Medical Science. M.W., E.H.K.S. and F.P. acknowledge funding by the iMOL (interfacing image analysis and molecular life-science) DFG Research group (project # 414985841, GRK 2566). F.P. acknowledges funding by the EU Horizon2020 project BRIGHTER (Grant #828931), the EU Horizon-EIC-2021 project B-BRIGHTER (Grant #101057894), and the German Space Agency at DLR (Grant #50W2019 and #50WB2316).

## Authors contribution

S.A.H., F.P. and M.W. designed the experiments. S.A.H. conducted most of the experiments described in the work. V.P. prepared the electrospun substrates and characterized them. F.P. and M.W. supervised the research. E.H.K.S. supported the research. The manuscript was written by S.A.H. and F.P. V.P. wrote the sections on the characterization of the electrospun substrate. All authors revised and agreed on the submitted manuscript.

## Declaration of interests

The authors declare no competing interests.

## Notes

### Competing Interest Statement

The authors have declared no competing interest.

